# VirRep: accurate identification of viral genomes from human gut metagenomic data via a hybrid language representation learning framework

**DOI:** 10.1101/2023.03.10.532047

**Authors:** Yanqi Dong, Wei-Hua Chen, Xing-Ming Zhao

## Abstract

Accurate identification of viral genomes from metagenomic data provides a broad avenue for studying viruses in the human gut. Here, we introduce VirRep, a novel virus identification method based on a hybrid language representation learning framework. VirRep employs a context-aware encoder and a composition-focused encoder to incorporate the learned knowledge and known biological insights to better describe the source of a DNA sequence. We benchmarked VirRep on multiple human gut virome datasets under different conditions and demonstrated significant superiority than state-of-the-art methods and even combinations of them. A comprehensive validation has also been conducted on real human gut metagenomes to show the great utility of VirRep in identifying high-quality viral genomes that are missed by other methods.

## Main

Viruses, especially bacteriophages (viruses that infect bacteria and archaea), are essential players of microbial communities and the most numerous biological entities within the microecosystem of human gut, having a fundamental impact on regulating the structure and function of microbial communities through widespread phage predation, lysogeny and horizontal gene transfer^1,2^. The gut virome has been heavily implicated in many human diseases, including inflammatory bowel disease^3,4^, type 2 diabetes^5,6^, and severe acute malnutrition^7^, to name just a few. Yet, our knowledge about the viral genomic diversity in the human gut increases at a slow pace for decades due to the difficulties in virus isolation. Metagenomic sequencing can produce huge amount of reads from prokaryotes (bacteria and archaea) as well as viruses in the microbial community regardless of cultivability. Identifying viral sequences from metagenomic data provides researchers another broad avenue for studying viruses in the human gut.

Many computational approaches have been designed to identify viruses from metagenomic data, which can be roughly grouped into two categories: alignment-based approaches and alignment-free approaches. Alignment-based approaches, such as VirSorter^8^, VIBRANT^9^, and VirSorter2^10^, discriminate between viral sequences and prokaryotic ones based on the combination of gene annotation results and genomic structural features. These methods typically start with gene prediction, then annotate the origin and function of the predicted proteins against a built-in database via multiple sequence alignment. The annotation results were compiled into quantifiable metrics, and finally along with other genomic structural features compared to a statistical null model or fed into a machine learning/deep learning model to determine the likelihood of a sequence being viral in origin. This set of methods is able to capture viral sequences with enough similarity to those in the reference database while keeping high specificity. However, their performance declines quickly when sequences are shorter than 10 kbp and the alignment procedure consumes too much time according to our evaluation. Moreover, high-quality reference genomes and well annotated genes are just the tip of the iceberg for viruses in human gut, severely ruining sensitivity of these methods to discover novel viruses.

Recently, alignment-free approaches, an alternative route for improved identification of viruses, have made profound progress to overcome the limitations of alignment-based approaches. Such methods are generally learning-based, leveraging machine learning or deep learning techniques to automatically learn the rules of k-mer usage hidden in viral and host genomes and utilize the differences to discriminate between them. For example, VirFinder^11^ trained three logistic regression models with lasso regularization on 8-mer frequencies to predict sequences in different length. DeepVirFinder^12^ and Seeker^13^ use the one-hot encoded sequence as input for Convolutional Neural Network and Long Short-Term Memory model, respectively. PPR-Meta^14^ encoded the sequence with one-hot embedding on both base and codon level, and constructed a Bi-path Convolutional Neural Network to simultaneously distinguish viruses, plasmids, and chromosomes. One special case is INHERIT^15^, which adopted the pre-train-fine-tune paradigm similar to that in natural language processing. In particular, INHERIT first pre-trained two BERT models in a self-supervised manner on virus and bacteria datasets respectively to learn the semantics and syntactic rules of DNA “words” (i.e., *k*-mers). It then added a simple classifier on top of the two pre-trained models and fine-tuned them simultaneously to distinguish between viruses and bacteria.

Despite the great value of these alignment-free approaches, directly applying them to human gut metagenomic data meets some challenges. First, from the modeling point of view, most existing methods just apply a simple neural network architecture. The shallow representations they produce are not competent to accurately distinguish viral sequences from prokaryotic ones, leading to much higher false positives compared to alignment-based approaches. This would be amplified to disastrous considering that the viral sequences make up a rather small proportion in typical human gut metagenomic samples. Although INHERIT, the exception, can obtain globally contextualized and informative representations of DNA sequences, it is computationally intensive and time consuming with a huge number of parameters approaching 200 million, thus not suitable for large-scale metagenomic data mining. Second, in terms of prediction process, none of the proposed methods consider viral sequences residing in host genomes (i.e., proviruses). Given the large part of temperate phages in human gut viral population, such ignorance would miss a significant group of viral sequences.

Here, we present VirRep, a novel method based on a hybrid DNA language representation learning framework for accurate and efficient identification of viral sequences from human gut metagenomic data. VirRep employs a context-aware encoder (Semantic Encoder) and a composition-focused encoder (Composition Encoder) to incorporate the learned knowledge and known records to better describe the source of a DNA sequence. Unlike existing methods, both the positive (virus) and negative (prokaryote) training sets are human gut centric. We also designed an iterative segment extension mechanism to extract viral signals from prokaryotic genomes. Benchmark results on mutiple datasets under different conditions show VirRep significantly outperforms state-of-the-art methods and even combinations of them on both bulk and virus-enriched metagenomic samples, especially for sequences shorter than 10 kbp. We also comprehensively validate the utility of VirRep to identify high-quality viral genomes that are missed by other methods from real human gut metagenomic samples.

## Results

### Overview of VirRep workflow and the designed representation learning framework

VirRep is a fully automated and end-to-end tool for accurately and efficiently mining viral signals from human gut metagenomic data based on sophisticated DNA language representation learning. VirRep consists of an input module and a scoring module (Fig. 1a). For each DNA segment in fixed length of 1 kbp and its reverse complementary strand, the input module first splits them into two non-overlapping 500 bp-long pieces, respectively. Each of the four pieces is then tokenized into a sequence of 7-mers, and a special token [CLS] is added in front of every 7-mers sequence. The scoring module is a deep siamese neural network consisting of a forward scoring system and a reverse scoring system. The two scoring systems separately assess the viral potential of the original sequence and its reverse complement, but share the same weights. The final output is defined as the average of the predictions from both strands.

**Fig. 1.**
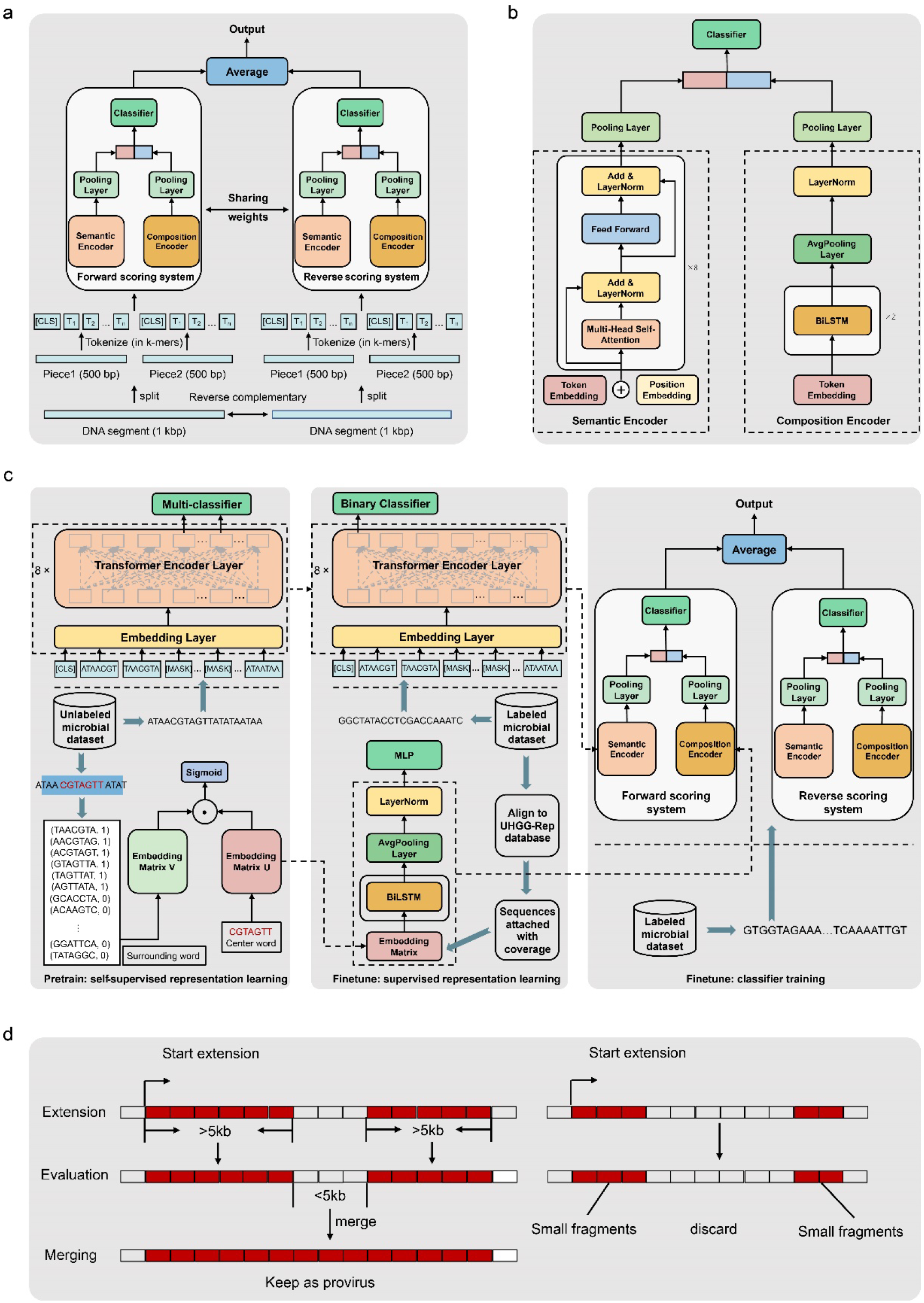
Schematic workflow of VirRep and overview of the hybrid DNA language representation learning framework. **a**, The input sequence is first split into several segments of 1 kbp in length. The original segment and its reverse complement are then preprocessed into 4 sequences of 7-mers, which are later fed into the two scoring systems. Final prediction is made by averaging the two scores. **b**, VirRep employs a siamese neural network with sharing weights, each composed of one Semantic Encoder, one Composition Encoder, two parallel pooling layers and a classifier. Semantic Encoder is a BERT-like neural network aimed to facilitate better global representation of the input sequence by obtaining fine-grained meaning of DNA words from the context. Composition Encdoer consists of an embedding layer, two BiLSTM layers, an average pooling layer and a layer normalization module, which is designed to encode alignment and similarity information of the input sequence against UHGG-Rep database. **c**, A hybrid training strategy is used to train VirRep. Semantic Encoder and the embedding layer are first pre-trained separately based on masked language model and skip-gram method, respectively. Then, Semantic Encoder is fine-tuned to predict whether a given sequence is of viral origin, and Composition Encoder to infer the proportion of the input sequence covered by UHGG-Rep database. Finally, two parallel pooling layers and a classifier are appended to the two encoders and fine-tuned simultaneously for virus identification. Dotted arrows represent model weights transfer. **d**, Iterative segment extension mechanism for virus identification and provirus extraction.

The main parts of the two scoring systems are Semantic Encoder and Composition Encoder, which are developed to take advantage of both alignment-based and alignment-free approaches (Fig. 2a). Specifically, Semantic Encoder employs a BERT-like^16^ architecture, relying on the multi-head self-attention mechanism^17^ to capture the importance of each DNA word (i.e., k-mer) and word-word dependencies. Composition Encoder is a stacked neural network composed of a token embedding layer, two BiLSTM^18^ layers, an average pooling layer and a layer normalization^19^ module. Composition Encoder behaves like a searching engine to retrieve alignment information of the input sequence against a latent database. Representations generated by the two encoders are non-linearly transformed by the two parallel pooling layers and then concatenated into a single tensor that incorporates globally contextual information and sequence alignment information. A simple classifier then outputs a value based on the informative tensor to measure the probability that the input sequence derives from viruses.

**Fig. 2.**
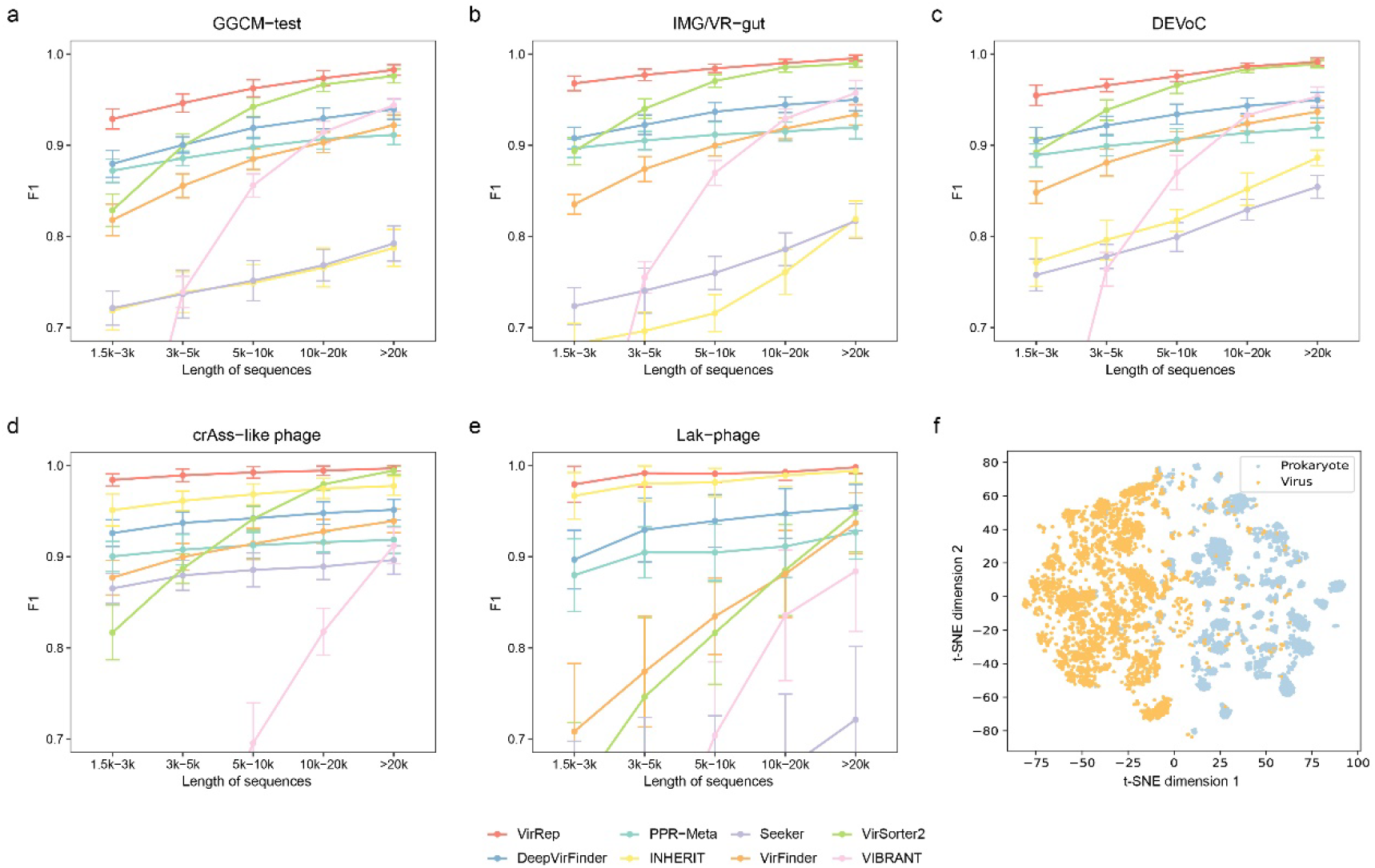
Performance evaluation of VirRep on multiple human gut virome datasets. **a-e**, VirRep consistently achieved better performance than existing methods on the **(a)** GGCM-test dataset, **(b)** IMGVR-gut dataset, **(c)** DEVoC dataset, **(d)** the set of crAss-like phages and **(e)** the set of Lak-phages across a broad range of sequence length. We conducted experiments multiple times on the five benchmarking datasets (*N* = 30 for GGCM-test, IMG/VR-gut, DEVoC and crAss-phages; *N* = 20 for Lak-phages). Each time an equal number of viral (*n* = 1000 for GGCM-test, IMG/VR-gut and DEVoC; *n* = 500 for crAss-phages; *n* = 100 for Lak-phages) and prokaryotic sequences were mixed. Error bars show the 95% confidence intervals. F1 score is used to measure the overall performance of VirRep and competing methods, while detailed recall and precision results are provided in Fig. S1 and Fig. S2. **(f)** The embedding generated by VirRep well separates viral and prokaryotic sequences. Shown is a two-dimensional visualization of the embedding vectors of 5000 randomly selected viral sequences from the GGCM-test dataset and 5000 randomly selected prokaryotic sequences from the IMG-GEM dataset based on t-SNE algorithm.

We adopted the pre-train-fine-tune paradigm to build VirRep on human gut centric datasets (Fig. 1c). We took advantage of both self-supervised and supervised manners to design a sophisticated DNA language representation learning framework. The two encoders were first separately pre-trained in a self-supervised manner. We used the adapted masked language model^20^ as the pretext task for Semantic Encoder pre-training to learn the semantics and syntactic rules of DNA words. While for Composition Encoder, we first pre-trained its embedding layer by predicting the surrounding contexts given the center word, similar to what has been done in the skip-gram model^21,22^. The aim of this step is to force similar 7-mers having similar embeddings. Next, we constructed two prediction tasks to finetune the two encoders to obtain fine-grained representations in a supervised manner. In particular, Semantic Encoder was made to predict whether the input sequence is of viral origin and Composition Encoder to infer the proportion (i.e., coverage) of the input sequence being covered by UHGG-Rep database. Finally, we simultaneously fine-tuned the two encoders with a classifier on top of the concatenated representation to build the final model for virus identification.

We also introduced a delicate mechanism for improved extraction of proviruses through iterative segment extension (Fig. 1d). For each sequence to be classified, we break it up into several 1 kbp-long segments. VirRep first assigns a score to every segment and takes the average as the score of the entire sequence. If the average score is above the cutoff set by the user, it is regarded significant and the sequence is considered as entirely viral. Otherwise, we search for segments with significant scores. We iteratively extend one segment at a time from the first one as long as the average score of these segments stay significant. The procedure is repeated until there is no significant segment left. We additionally provide a filter to prevent accidental false positives. An initial extracted subregion is retained only if it is longer than the user-set minimal length (5 kbp in defualt) or a given percentage (0.5 in default) of the length of the original sequence. To enhance the continuity of the hits, the filtered subregions deriving from the same sequence are merged if the gap between them is shorter than the pre-defined threshold (5 kbp in defualt) or the maximum proportion (0.5 in default) of their summed length.

### VirRep enables a robust detection of viruses on multiple datasets across a broad range of sequence length

We first evaluated the newly proposed method VirRep on multiple recently published human gut virome datasets at different sequence lengths. These datasets included the test set of the combination of GVD^23^, GPD^24^, CHVD^25^ and MGV^26^ (referred to GGCM-test), subset marked as human intestinal origin of the virus genome database IMG/VR v3 (ref. ^27^) (referred to IMG/VR-gut), the Danish Enteric Virome Catalog^28^ (referred to DEVoC), the union set of complete and high-quality crAss-like phages from two studies^29,30^ and a set of largest reported megaphages to date in the human gut microbiome^31^ (referred to Lak-phage). As negative control, prokaryotic sequences assembled from human gut metagenomic samples were also included. In order to evaluate the performance of VirRep at different sequence lengths, several fragments were randomly generated for a given length interval from each qualified original genome in the test datasets. We compared VirRep against both popular alignment-based (VIBRANT and VirSorter2) and alignment-free (VirFinder, DeepVirFinder, PPR-Meta, Seeker and INHERIT) approaches.

The overall performance of each method was first assessed on the GGCM-test dataset to see how sequence length will affect their performance. Sequences in this dataset did not overlap with those in the training set and shared less than 90% nucleotide identity over 80% of their length with the training ones. As shown in Fig. 2a, all methods demonstrate an increased accuracy as sequence length grows, while VirRep significantly outperforms the other methods with the highest average F1 score and lowest standard error at all length intervals. Specifically, VirRep achieved F1 score of 0.93, 0.95, and 0.96 for sequences of 1.5-3 kbp, 3-5 kbp, and 5-10 kbp, while the corresponding values for the second best result were 0.88, 0.90, and 0.94, reflecting 5.7%, 5.6%, and 2.1% increase, respectively. Although VirSorter2, the top alignment-based method, were comparable with VirRep for sequences longer than 10 kbp, its performance declined sharply with shorter sequences, especially for those no more than 10 kbp. Several alignment-free approaches, such as DeepVirFinder and PPR-Meta obtained relatively decent results with sequences of length <5 kbp, their enhancement, however, was not as good as expected when sequences reached 10 kbp and longer. Seeker and INHERIT performed poorly on this dataset with F1 score <0.8 at all length intervals. These results were further validated on the IMG/VR-gut and DEVoC datasets (Fig. 2b, c). The two datasets were generated mainly by explicit homology searching to known viruses and showed higher sequence similarity with our training set, hence explaining the performance improvement achieved by all the evaluated methods.

We additionally evaluated VirRep and the other methods on two particular viral clades in the human gut: crAss-like phages and Lak-phages. CrAss-like phages represent the most abundant and prevalent viral family in human gut microbiota^29,30,32^. Effective identification of their genome sequences from the human gut metagenomes is of great importance for phage biology study and function characterization. Nearly all the methods performed well at most length intervals, of which VirRep showed the best result, with F1 score exceeding 0.98 even when sequences were as short as 1.5-3 kbp (Fig. 2d). VirSorter2, typically the second-best method on the three previously described datasets, had much lower F1 scores (decreased by 6.4-19.9%) than VirRep for sequences shorter than 10 kbp. INHERIT, nevertheless, replaced other methods as the most excellent tool besides VirRep unexpectedly. Lak-phages, the largest phages ever reported in human gut microbiome with genome size >540 kbp, are estimated to infect *Prevotella* and be widespread in the population consuming non-Western (i.e., high-fiber and low-fat) diets. Similar trends were also observed on this test set (Fig. 2e). Alignment-free methods (except for Seeker) generally obtained higher F1 scores than the two alignment-based tools, and VirRep still held the top position. Interestingly, the performance of VirSorter2 dramatically decreased by 5.0-35.0% compared to VirRep at the five length intervals, indicating its limitation to identify novel and previously overlooked viruses in the human gut. We noticed that INHERIT performed much better on these two datasets, likely due to the overlap between them and the training set of INHERIT.

In order to intuitively explore the ability of VirRep to discriminate between viruses and prokaryotes, we visualized the sequence embeddings using t-SNE^33^. As shown in Fig. 2f, most of the sequences were well separated according to their origin even in a two-dimensional space. Overall, VirRep is the only method that enables accurate identification of viruses on all the tested datasets across a broad range of sequence length.

### Dedicated encoders and representation learning improve sensitivity and specificity of virus identification

To further explore what roles of the two encoders are playing and the impact of the designed representation learning, we conducted two sets of ablation experiments on the GGCM-test dataset. We first compared the performance among the full implementation of VirRep, the fine-tuned Semantic-Encoder-based classifier and the finetuned Composition-Encoder-based predictor from various aspects. The results showed that VirRep significantly outperformed the other two components on precision, F1 score and true negative rate (TNR), and displayed comparable results with the Semantic-Encoder-based classifier on recall (Fig. 3a). The enhancement was particularly pronounced for sequences shorter than 10 kbp. Moreover, the performance of the Semantic-Encoder-based classifier was obviously better than that of the Composition-Encoder-based predictor in most cases, which reveals that the representation generated by Semantic Encoder is the primary foundation for determining the label of the input sequence. Besides, we also noticed that the Composition-Encoder-based predictor tended to relatively better identifying prokaryotic sequences than viral ones, with its true negative rate 3.7-10.3% higher than the corresponding recall value. Such observation indicates that Composition Encoder acts as an assistant consultant to correct the mistakes made by Semantic Encoder, which is consistent with our hypothesis that the alignment information will facilitate the improvement of virus detection precision. All in all, the two dedicated encoders together can effectively combine the learned knowledge and the known records to better describe a sequence, thus enabling more precise and sensitive identification of viral genomes.

**Fig. 3.**
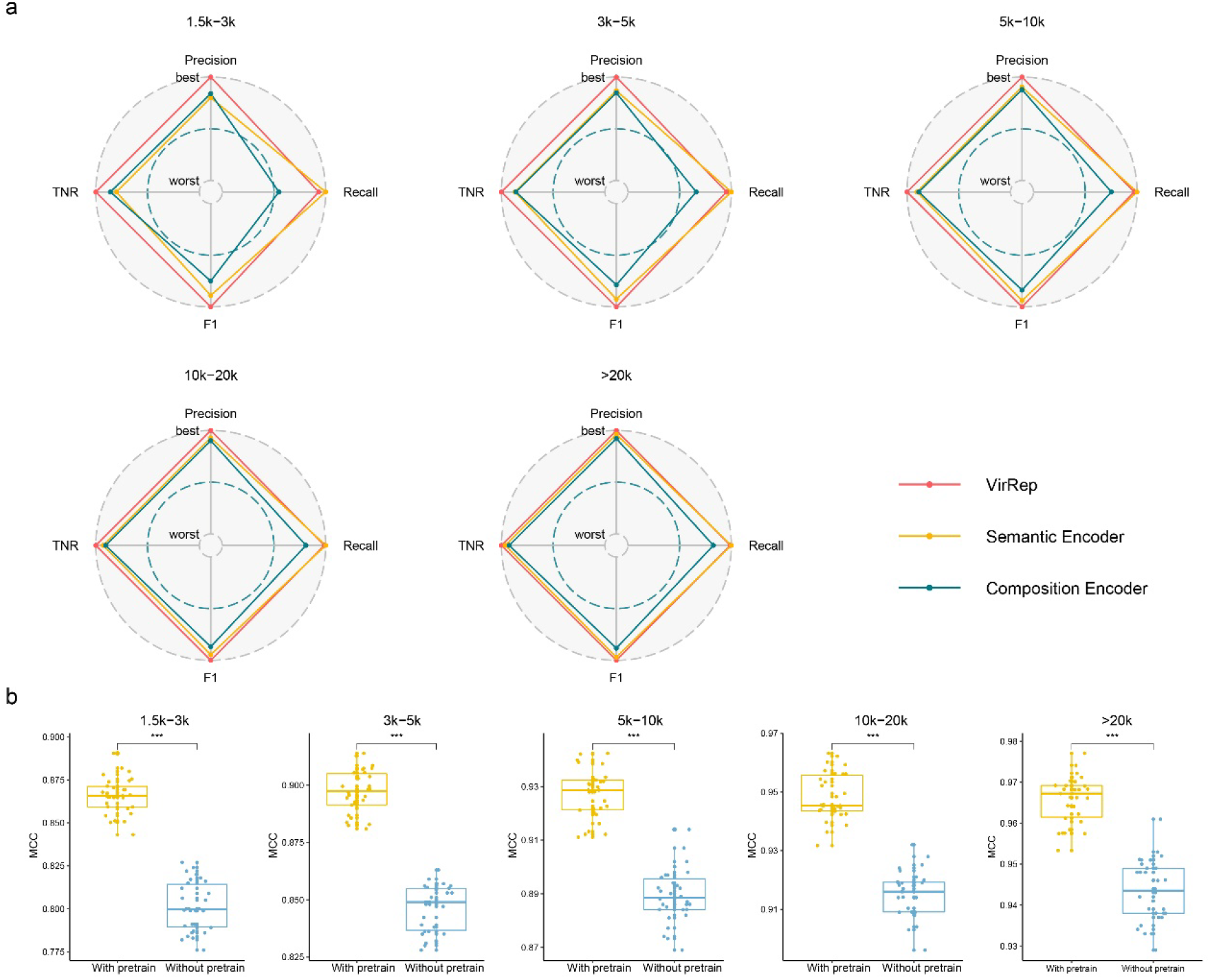
The two encoders of VirRep and pre-training significantly improve the overall performance of virus identification. Shown are the ablation experiments on the GGCM-test dataset under different sequence length intervals. Experimental results on the IMG/VR-gut dataset are available in Fig. S3 and Fig. S4. **a**, Radar plots showing the relative superiority on precision, recall, F1 score and true negative rate (TNR) among the full implementation of VirRep, the Semantic-Encoder-based classifier and the Composition-Encoder-based predictor. The values are linearly scaled for better visualization. **b**, Boxplots showing the distributions of Matthews correlation coefficient (MCC) of the model with pre-training (With pretrain) and the model training from scratch (Without pretrain) over 30 replicates. Hypothesis tests were performed based on two-sided Wilcoxon signed-rank test.

Pre-training can often help the model learn common and basic knowledge from huge amount of unlabeled data. In order to explore whether the learned knowledge would be beneficial to our ultimate goal and what extent it would influence the model performance, we retrained the model from scratch and compared the retrained version to VirRep. Matthews correlation coefficient (MCC), which simultaneously takes true positives, false positives, true negatives and false negatives into account, is leveraged to measure the overall performance of the two models. The value of MCC ranges from -1 to +1, where a higher coefficient represents closer consistency between the observed and the predicted labels. As shown in Fig. 3b, VirRep with pre-training had evidently higher MCC than that of the version training from scratch across all the five sequence length intervals. The differences were not only significant in statistics (Wilcoxon signed-rank test, *P* = 1.86 × 10^−9^ for length interval 1.5k-3k, 3k-5k, 5k-10k and 10k-20k; *P* = 1.8 × 10^−6^ for length interval >20k), but also numerically distinct. The mean values of MCC over 30 replicates were 0.86, 0.90, 0.93, 0.95 and 0.97 as sequence length grows for VirRep with pre-training, while the corresponding values for the retrained version were 0.80, 0.85, 0.89, 0.91, and 0.94, reflecting 8.1%, 6.0%, 4.3%, 4.0% and 2.4% improvement, respectively. Such inspiring observations provide ample evidence on the tremendous benefits of pre-training in improving model performance on down-stream tasks.

In conclusion, the two dedicated encoders can well incorporate the learned knowledge and known records to better describe the source of a DNA sequence, thus enabling VirRep to take advantage of both alignment-free and alignment-based methods for more sensitive and precise viral signal mining from metagenomic samples. Pre-training facilitates the precious knowledge learned from unlabeled data to guide model training on down-stream task-specific data. The two encoders and representation learning together profoundly enhance the overall performance of virus identification.

### VirRep is well applicable to both bulk and VLP human gut metagenomic samples

Traditionally, the amount of viral DNA only accounts for a small (∼5.8% on average) part of the total DNA of the whole microbial community (bulk metagenomic samples) in human gut^34^. Although VLP metagenomic sequencing can enrich viruses to a certain extent, there is still unneglectable background noise from prokaryotic organisms^35^. We hence reasoned that the conventional test with equal number of viral and prokaryotic sequences may not faithfully reflect the performance of evaluated methods in real-world usage scenarios. To demonstrate the effectiveness of VirRep on both bulk and VLP human gut metagenomic samples, we focused our efforts on four simulated metagenomic datasets with different viral proportions.

We first assessed VirRep and other methods to see how precision and recall will vary with increasing thresholds (Fig. 4a). Here, we excluded the two alignment-based approaches, VIBRANT and VirSorter2, since their outputs attached no scores or merely assigned scores for significant hits. For the two mimic bulk metagenomic samples (viral proportion 5% and viral proportion 10%), VirRep was the only method allowing for decent detection of viral sequences, as the average AUPRC scores all exceeding 0.90, and were 7.1% and 4.4% higher than the second best results, respectively. This indicates VirRep is significantly more powerful than existing methods on bulk metagenomic samples. We also tested VirRep on two datasets with viral sequences made up equal or more than half of the whole community. These datasets were built to simulate virus-enriched metagenomes and the extreme case where VirRep is leveraged to decontaminate the virome samples. Although most of the assessed methods performed well on these two cases, VirRep still achieved the highest AUPRC scores and had the highest precision at any recall value. Together, VirRep can be applied to both bulk and VLP human gut metagenomic samples and is the most powerful method across a broad range of viral proportions.

**Fig. 4.**
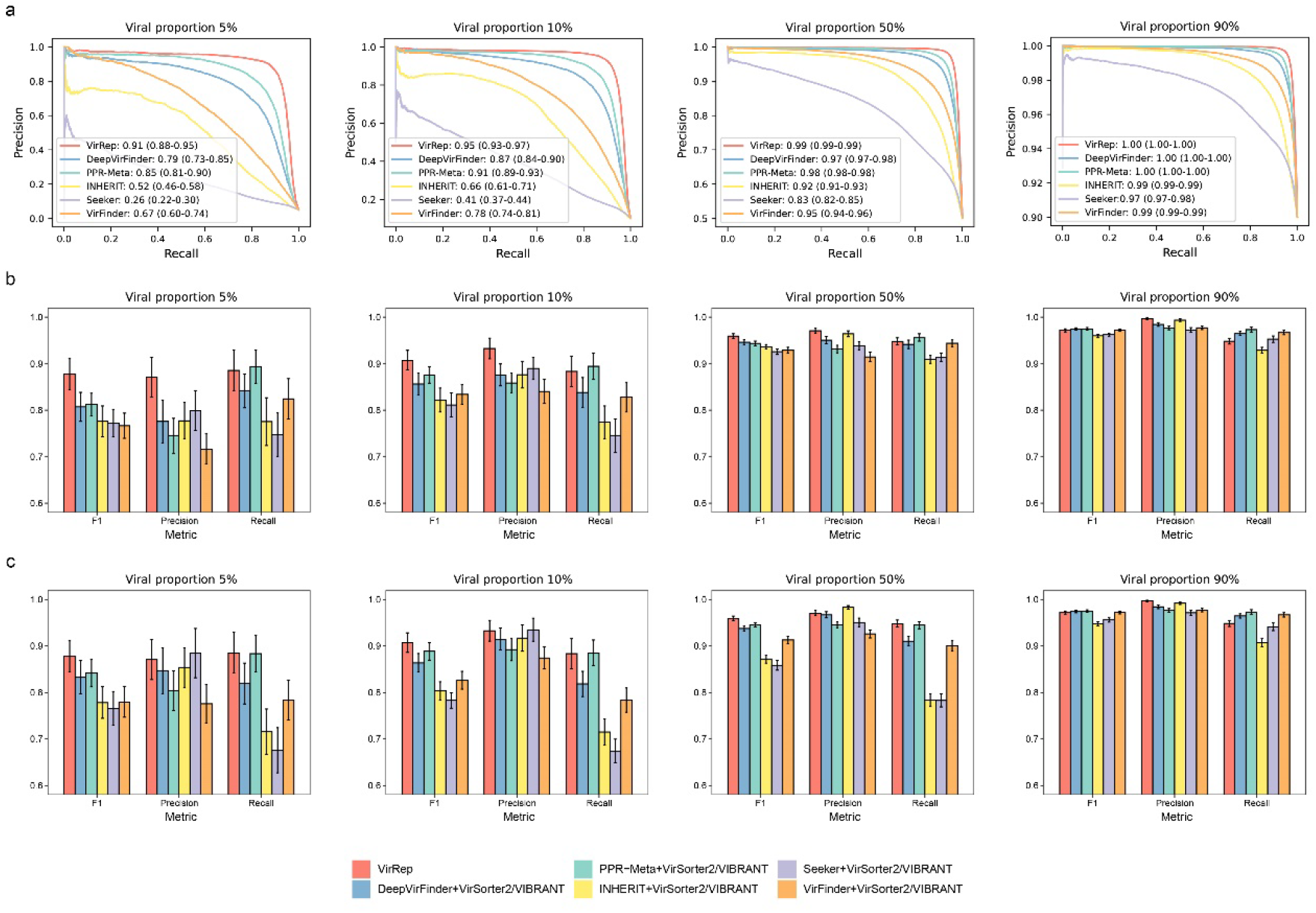
Comparison of the performance of VirRep against those of other methods and pipelines on simulated metagenomic samples with different viral proportions. Shown are the results on samples constructed based on the GGCM-test dataset. Results on samples constructed based on the IMG/VR-gut dataset can be accessed in Fig. S5-7. **a**, Precision-recall (PR) curves and AUPRC values for VirRep and the five alignment-free methods under viral proportion 5%, 10%, 50% and 90%, respectively. Values in the bracket represents the lower bound and upper bound of the 95% confidence intervals over 30 replicates (5000 sequences in total for each replicate). **b**, Average F1 score, precision and recall for VirRep and pipelines composed of VirSorter2 and one alignment-free method under viral proportion 5%, 10%, 50% and 90%. **c**, Average F1 score, precision and recall for VirRep and pipelines composed of VIBRANT and one alignment-free method under viral proportion 5%, 10%, 50% and 90%. Error bar shows the 95% confidence intervals over 30 replicates.

Alignment-based and Alignment-free approaches each can handle different sequence space, and are usually complementary. Combining the two categories of methods would be a promising means for virus identification. Many studies have applied this strategy to mine viral signals from metagenomic samples^23,24^. We then compared VirRep against several mixed virus identification pipelines each consisting of one alignment-based method and one alignment-free method. Surprisingly, VirRep alone obtained better or comparable results than the complicated pipelines. The superiority was especially marked when prokaryotic sequences dominate the samples (Fig. 4b). For example, the F1 score of VirRep were 0.88 and 0.91 for datasets with 5% and 10% viral sequences, which is 8.1% and 3.6% higher than the best pipeline contained VirSorter2 (i.e., VirSorter2+PPR-Meta), respectively. This was induced by the stronger capacity of VirRep to differentiate between viruses and prokaryotes, since VirRep had much higher precision (16.9% and 8.7% higher on average) while keeping similar sensitivity compared with the pipeline composed of VirSorter2 and PPR-Meta. For the virus-enriched metagenomes, VirRep still achieved the best overall performance, while DeepVirFinder replaced PPR-Meta to be the best partner with VirSorter2. Even in the extreme case where viral sequences accounts for the vast majority of the samples, VirRep remained on par with the state-of-the-art strategy. Similar trend was also observed when comparing against the pipelines with the participation of VIBRANT. VirRep achieved significantly higher precision and recall than most of the pipelines and topped the list in overall performance (F1 score) on datasets with 5%, 10% and 50% viral sequences, and were comparable with the tested pipelines when viral sequences made up the majority of the metagenomic sample (Fig. 4c).

In summary, VirRep is not only well suited for both bulk and VLP human gut metagenomic samples, but also can achieve better performance than existing methods or even a combination of them in most cases.

### Identify high-quality viral genomes from real human gut metagenomic samples of a CRC cohort using VirRep

Encouraged by the above benchmark results, we applied VirRep to scan the real human gut metagenomes of a colorectal cancer (CRC) cohort with 75 CRC patients and 53 healthy controls. In this section, we tended to practically demonstrate the utility of VirRep to identify high-quality viral genomes that may be missed by other methods.

Sequencing reads from each of the 128 metagenomes were first preprocessed to filter out human-genome-derived reads and low-quality bases, then assembled by MEGAHIT. The resulting contigs were screened by VirRep and other methods (including VIBRANT, VirSorter2, VirFinder, DeepVirFinder, PPR-Meta, Seeker and INHERIT) to identify putative viral sequences. Since we focused on high-quality viral genomes, only hits longer than 5 kbp were kept for further analysis. After removing flanking host regions and dereplication, we obtained a collection comprised of 15519 viral representative genomes using VirRep (Methods).

CheckV was leveraged to estimate the level of completeness of each viral genome. In total, 501 genomes were predicted as complete and 1444 as high quality (>90% complete) (Fig. S8). Among these complete and high-quality viral genomes, 130 were totally ignored or partially detected as genome fragments (<90% completeness) by other methods. We explored in detail six of these complete and high-quality viral genomes with genes encoding termianse large subunit. The first genome (ERR1018185_k119_210575) has 55,997 bp and is estimated to be approximately 96% complete based on amino acid identity (AAI) with high-confidence. We identified 76 protein-coding genes and annotated 25 (∼33%) of them, including the three hallmark genes in tailed phages: large terminase subunit, major capsid protein and portal protein (Fig. 5a). The second genome (ERR1018270_k119_58275) is in length of 42,488 bp and determined to be 100% complete with 39 predicted genes, of which we were able to annotate 16 (∼41%) of them (Fig. 5b). Two of the annotated genes encode structural proteins (i.e., large terminase subunit and major capsid protein). More interestingly, the virus is predicted to encode holin, showing its potential to specifically control pathogenetic bacteria. The third genome (ERR1018186_k119_78063) is 34,816 bp-long with a lower level of completeness (∼92%). It was predicted to encode 36 genes and 11 (∼31%) of them could be annotated, again including terminase large subunit, major capsid protein and portal protein (Fig. S9). Phylogenetic analysis was performed based on the large terminase genes and showed the three viruses are all members of the *Siphoviridae* family (Fig. 5d). The fourth genome (ERR1018188_k119_117010) has 36,519 bp with an estimated completeness of 97%. We recognized 52 putative genes and annotated more than half (31, ∼60%) of them (Fig. S10). The fifth genome (ERR1018262_k119_108237) has 39,369 bp and is predicted 100% complete with 51 putative genes, of which we annotated 28 (∼55%) of them (Fig. 5c). We found one gene encode endolysin in each of the two viral genomes, indicating both the two phages can be of great value in medical treatment. We build phylogenetic trees relying on the large terminase subunits and inferred the two viruses belong to the *Myoviridae* family (Fig. 5e). The last genome (ERR1018304_k119_47942) we analyzed is 33,711 bp-long and 94% complete with 36 predicted genes, of which 23 (∼64%) were annotated, including an integrase-encoding gene, which indicates the genome may derive from a prophage (Fig. S11). Phylogenetic analysis of the large terminase subunits demonstrated the virus is a distinct member of the *Podoviridae* famlily (Fig. 5f).

**Fig. 5.**
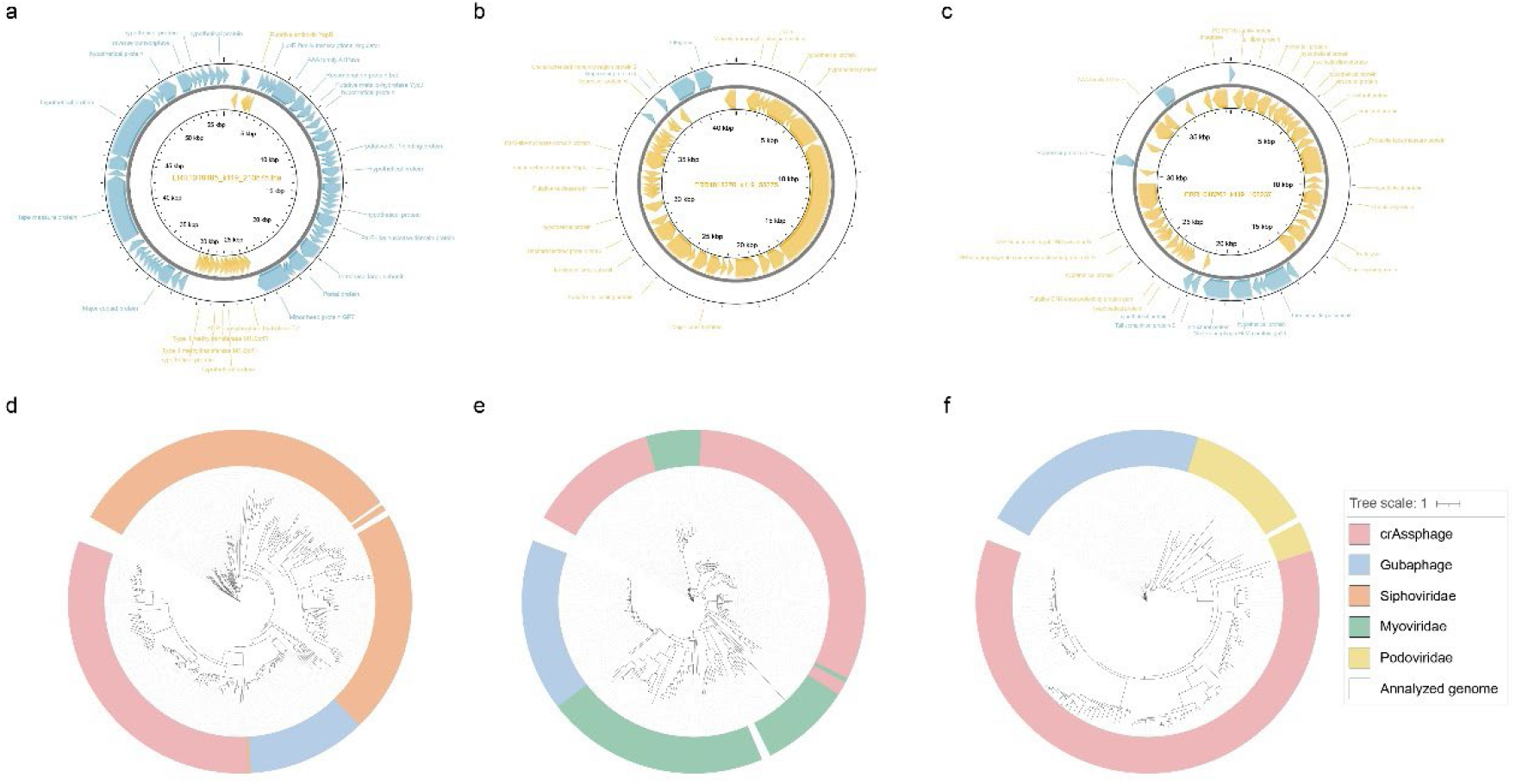
Example of high-quality genomes identified by VirRep and missed by other methods. **a-c**, Annotated gene map of the viral genomes: **(a)** ERR1018185_k119_210575, **(b)** ERR1018270_k119_58275, and **(c)** ERR1018262_k119_108237. **d-f**, Phylogenetic trees of large terminase subunits for the six viral genomes: **(d)** ERR1018185_k119_210575, ERR1018270_k119_58275 and ERR1018186_k119_78063; **(e)** ERR1018188_k119_117010 and ERR1018262_k119_108237; **(f)** ERR1018304_k119_47942.

## Discussion

Viruses are often recognized as the ‘dark matter’ of the human gut microbiome, profoundly impacting the human health through microbial community regulation. Accurate and efficient identification of viruses from the mixed pool of metagenomic sequences has been the cornerstone of human gut virome studies.

In this article, we present a novel method, named VirRep, to identify viral sequences from human gut metagenomic data. The core modules of VirRep lie on Semantic Encoder and Composition Encoder, which are designed to take advantage of alignment-free and alignment-based methods, respectively. Ablation studies demonstrate that the two encoders can well incorporate the learned knowledge and known records to better characterize DNA sequence. We also propose a hybrid DNA language representation learning framework that combines self-supervised and supervised manners to obtain fine-grained representations for predicting the origins of input sequences. Unlike existing alignment-free approaches, we trained VirRep on large-scale human gut centric microbiome datasets, thus enabling VirRep to better learn the specific sequence patterns of viruses and prokaryotes residing in human gut. VirRep displays high accuracy and robustness on multiple human gut virome datasets across a broad range of sequence length (Fig. 2a-e), and shows significant superiority or comparable performance on both bulk and VLP metagenomic samples comparing against existing methods and even the combination of them (Fig. 4). We additionally demonstrate the utility of VirRep by applying the method to a CRC cohort with 128 human gut metagenomic samples, where we identified several high-quality viral genomes associated with CRC that were missed or partially detected by other methods (Fig. 5).

Despite the inspiring results, there are also some limitations and promising directions for further improvement should be noticed. First, VirRep is now dedicated to human gut metagenomic samples. The performance of identifying viruses from other environmental samples have not been comprehensively tested and are not guaranteed, therefore directly applying VirRep to other habitats, such as soil and ocean, needs to be cautious. In the future, we will consider to pre-train the model on pooled datasets from all habitats to learn more general knowledge and transfer the pre-trained model to various environments. Second, the size of Semantic Encoder is reduced to be much smaller compared to the original version of the BERT-base model to compensate for running efficiency, which will inevitably make a compromise on the model performance. Knowledge distillation^36^ in the form of teacher-student framework can be taken into account for better model compression. Finally, the skip-gram method used to pre-train the embedding matrix of Composition Encoder is indirect for our purpose and does not consider the effect of the positions of differential bases in the central k-mer on surrounding words in the context window. Take the two 7-mers with only one differential base as an example, they can have up to 5 identical surrounding words for window size set to 5 if the differential base appears at the head or the tail of the k-mers, while merely 4 identical surrounding words at most when the differential base occurs at other locations. A more direct metric measuring the distance between two k-mers holds great potential to further improve the pre-training process.

## Methods

### Neural network architecture of VirRep

We employ a siamese neural network to separately score the original sequence and its reverse complementary strand. The siamese network has two sub-networks with the same architecture and shares the same weights. Each of the two sub-networks contains one Semantic Encoder and one Composition Encoder, which are designed to take advantage of alignment-free and alignment-based approaches, respectively. Apart from the two encoders, there are also a pooling module and a binary classification layer on top of each sub-network.

#### Semantic Encoder

Semantic Encoder is generally a BERT-like^16^ neural network but with a reduced size (see Supplementary …) to enhance running efficiency for easy access to large-scale metagenomic data mining. The structure of Semantic Encoder consists of an embedding layer and eight Transformer encoder blocks^17^. The embedding layer comprises two lookup tables, which stores the embedding vectors of k-mers and positions respectively. Each Transformer encoder has two sub-layers. One is the multi-head self-attention mechanism, and the other a position-wise feed-forward neural network. The two sub-layers both has a residual connection around, and is followed by layer normalization operation. The core of Transformer encoder is the multi-head self-attention mechanism, which can be formular as follows:

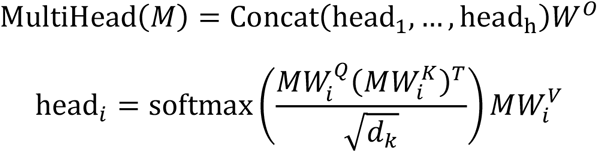

where *W*^*O*^ and 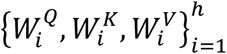 are the learned parameters, and *M* is the embedding matrix representing the input sequence for the first block and the output of the last block for others.

A tokenized DNA sequence was first fed into the embedding layer and transformed into two matrices encoding words and word positions independently. The two matrices are then added and used as input to the Transformer encoders. The multi-head self-attention mechanism adjusts the embeddings of each word according to the contexts. And the output of the last Transformer encoder block corresponding to the first token [CLS] is extracted as the final semantic representation of the entire sequence. The data stream can be expressed as:

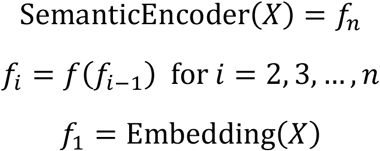

where *X* represents the one-hot encoding matrix of the input sequence, and *f* the function underlying Transformer encoder.

#### Composition Encoder

Composition Encoder is designed to encode the base order and content of a sequence for fast retrieval from the latent prokaryotic database and rough similarity estimation. Little contextual understanding of the sequence is required for such task, but the k-mer itself and its location in the sequence count. Hence, LSTM would be a fit choice to achieve this goal.

Concretely, Composition Encoder is composed of an embedding layer, two stacked BiLSTM layers, an average pooling layer and a layer normalization module. The embedding layer converts the tokenized DNA sequence into a matrix, which is then fed into the BiLSTM blocks. The resulting output of each token incorporates its forward and backward information about sequence composition and can be regarded as a candidate embedding of the entire sequence. Average pooling is applied to reduce the dimension by taking average on the outputs of all the tokens. After layer normalization, the final output is treated as the compositional representation of the input sequence.

In formular, the process of feeding a DNA sequence *X* into Composition Encoder can be summarized as follows:

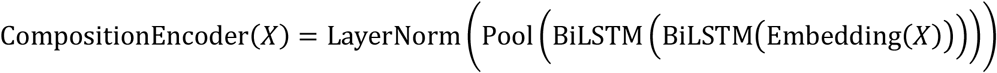

#### Pooling module and classification layer

There is a pooling layer above each of the two encoders for nonlinear transformation of the produced representation. The pooling layer is actually a fully connected feed-forward network with hyperbolic tangent activation function. The classification layer aims to output the likelihood of the input sequence deriving from viruses, which consists of a multilayer perceptron with a single hidden layer.

### Hybrid DNA language representation learning framework

We proposed a hybrid DNA language representation learning framework by adopting both self-supervised and supervised manners to better describe a sequence. The whole training process can be divided into three stages: pre-training, fine-tuning phase1 and fine-tuning phase2.

#### Pre-training

Pre-training aims to make the model learn the basic and common knowledge from huge amount of unlabeled data, which will aid to improve model performance on down-stream tasks. The key to pre-training lies on appropriate pretext tasks. We pre-trained Semantic Encoder and Composition Encoder independently and applied different pretext tasks according to their down-stream tasks.

##### Semantic Encoder pre-training

Semantic Encoder is pre-trained to acquire biological meaning and permutation rules of DNA words. Following the previous works^16,20^, we used an adapted version of masked language model as the pretext task. All DNA segments used for training are in fixed length of 500 bp. For each segment, we first tokenized it into a sequence of k-mers and inserted a special token [CLS] (representing the entire segment) at the head of the sequence. Here, we chose *k* = 7 to balance the model size and performance. We next uniformly selected 2.5% of the input tokens as the anchor, and extended to cover the most adjacent *k* − 1 tokens. The tokens and their extensions (15% of the input sequence) were selected for possible replacement, of which 80% were replaced with the [MASK] token, 10% kept unchanged and the remaining substituted by a random token. In the process of pre-training, we predicted what the masked token it was by feeding the final output of each masked token into a multi-class classification layer. The objective function is to minimize the cross entropy loss between the predicted likelihood and the true label. We optimized Semantic Encoder with AdamW^37^. More detailed settings of training are provided in Supplementary methods.

##### Composition Encoder pre-training

Unlike Semantic Encoder, only the embedding layer of Composition Encoder is pre-trained. To encode the compositional pattern of a sequence for retrieval and similarity estimation against a latent database, embedding the k-mers into a space where k-mers with high identity are closer is the fundamental step. We followed the skip-gram^21^ method to pre-train the embedding matrix of k-mers (*k* = 7), which could be briefly summarized as predicting the surrounding k-mers given a center k-mer. The rationality for choosing this pretext task is that similar center k-mers are more likely to have identical neighbors.

We applied the negative sampling technique during training^22^. Specifically, we first converted the nucleotide sequence into a sentence of 7-mers and set the context window size to 5. For a given center 7-mer, its neighbors in the context window form positive word pairs with it, respectively. For each positive word pair, ten negative word pairs were generated by randomly sampling other 7-mers not in the context window. The other 7-mer in the word pair is called target word for convenience. We maintained two embedding matrices *U* and *V* with the same size. Center 7-mers were fed into the matrix *U*, while the target 7-mers were fed into the matrix *V*. Then, we performed inner product on the embedding vector of center 7-mer and the embedding vector of each corresponding target 7-mer. And the scaler was finally normalized to be between 0 and 1 by sigmoid function. We used the binary cross entropy loss as the cost function and optimized the two embedding matrices by Adam^38^ algorithm (learning rate set to 0.001) with a batch size of 1000. After xxx training steps, the matrix *U* was kept as the initialized parameters for the embedding layer of Composition Encoder.

#### Fine-tuning phase1

After pre-training, we continued to fine-tuned the two encoders separately in a supervised manner on task-specific data to obtain fine-grained sequence representations. Rather than starting from randomly initialized model weights, we transferred the pre-trained parameters as the initialization before fine-tuning the network. The same training tricks were utilized for both encoders, where the learning rate was first warm up to a peak value and then linearly decayed.

##### Semantic Encoder fine-tuning

For Semantic Encoder, we forced the model to learn how to determine if a given nucleotide sequence derives from viruses, a sequence-level binary classification task. As mentioned above, the vector corresponding to the token [CLS] in the final output matrix (output of the last Transformer encoder block) is regarded as the aggregate representation of the entire sequence. We feed the vector into a classification layer which is a simple fully connected feed-forward neural network with the sigmoid activation function. Considering the fact that DNA sequence has double strands, the prediction scores should be identical for both strands. Thus, we score the original sequence and its reverse complement independently and take their average as the final output. We followed the instructions in Sun et al. (ref. ^39^) to fine-tune Semantic Encoder, including layer-wise decreasing learning rate and applying conservative learning rate to avoid catastrophic knowledge forgetting. For training details and hyperparameter settings, please refer to Supplementary Methods.

##### Composition Encoder fine-tuning

Composition Encoder was fine-tuned to infer the proportion of the input sequence covered by UHGG database, which can be viewed as a regression task. Unlike the pre-training process in which only the embedding matrix was trained, all components of Composition Encoder and an additional regression layer consisting of two fully connected networks were trained simultaneously at this time. We generated the labels of training data by aligning the sequences against UHGG database. Specifically, the viral sequences were directly blasted against UHGG database, while for prokaryotic ones, extra work should be introduced to ensure data diversity since the original training sequences themselves are part of UHGG database. We randomly selected half of the sequences in the prokaryotic training set and changed 15% the bases by deletion, insertion or substitution operations. Then, we blasted both the changed and unchanged prokaryotic sequences against UHGG database to carry out pairwise comparisons. We kept hits with E-value≤1e-8. For each query sequence, we merged the aligned regions that were ≥ 500 bp and shared at least 90% nucleotide identity with bedtools v2.29.1 (ref. ^40^). The coverage for each 500 bp-long segment was obtained by calculating the ratio of the length of the aligned regions to the length of the segment. We dealt with the double strands in the same way as Semantic Encoder. The cost function is Hubber loss, which is formalized as:

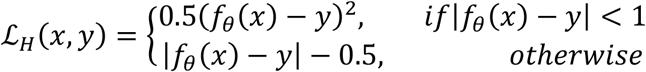

where *x* and *y* represents the input sequence and the attached coverage, respectively. *f*_*θ*_(·) denotes the function of Composition Encoder regressor and *θ* is the set of the learned parameters. The more detailed hyperparameter settings can be found in Supplementary Methods.

#### Fine-tuning phase2

The two encoders are now able to generate meaningful representations after fine-tuning separately on their own down-stream tasks. We then fine-tuned the pooling module and the classification layer over the two representations to discriminate between viruses and prokaryotes. The parameters of the two encoders were frozen at this step. Again, we employed binary cross entropy loss to measure the distance between the predicted probability and the true label. See Supplementary for more details of the final fine-tuning process.

### Applying VirRep to real human gut metagenomes

Raw sequencing reads of the 128 human gut metagenomic samples of a Chinese CRC cohort were downloaded from the NCBI SRA database under accession PRJEB10878. FastQC v0.11.9 (https://www.bioinformatics.babraham.ac.uk/projects/fastqc/) was first used to check the overall quality of each downloaded sequence data. We then removed human-derived reads by mapping them to human reference genome (hg38) using Bowtie2 v2.2.3 (ref. ^41^). Next, fastp v0.20.0 (ref. ^42^) was utilized to trim adapters and low-quality bases with the following parameters: ‘-l 50 -x -q 20 -u 5 -M 20 -W 4’. After preprocessing, MEGAHIT v1.2.8 (ref. ^43^) was used to assemble the high-quality clean reads into contigs with default parameters for each sample.

To identify viral sequences from the assemblies with high confidence, VirRep was run with parameters: ‘--min-score 0.8, --provirus-minlen 5000’, while VirSorter2 was run with parameters: ‘--high-confidence-only’ and VIBRANT with default parameters. As for the alignment-free methods, we ran the re-trained models independently except for INHERIT, which was run with the provided version. Sequences with score > 0.9 were considered as viral for the alignment-free methods. Since we focused on high quality genomes, only hits longer than 5 kbp were kept. CheckV^44^ was run on the kept sequences for the first time to remove potential flanking regions. The resulting cleaned viral hits were dereplicated by clustering these sequences at a 95% nucleotide identity over a local alignment of 85% of the shortest sequence using CD-HIT v4.8.1 (options ‘-c 0.95 -G 0 -aS 0.85’). Finally, we ran CheckV again on the non-redundant viral sequence set to assess the level of completeness of each genome.

### Gene prediction and functional annotation

Protein-coding genes were identified by prodigal-gv v2.10.0 (https://github.com/apcamargo/prodigal-gv) in metagenome mode (option ‘-p meta’), a fork of Prodigal^45^ meant to improve gene calling for giant viruses and viruses that use alternative genetic codes. Proteins translated from the CDS regions were then annotated with eggNOG mapper v1.0.3 (options ‘--Z 29033, --hmm_evalue 1e-5’) against VOGDB (https://vogdb.org/).

### Phylogenetic analysis of the six high-quality viral genomes

We downloaded the large terminase subunits of common viral families (such as *Siphoviridae, Myoviridae* and *Podoviridae*) from the NCBI viral RefSeq database^46^. Genomes for crAss-like phages were pooled from two studies as mentioned above and Gubaphages were obtained from GPD. The large terminase subunits of these two viral clades were extracted according to the gene functional annotation results. Phylogenetic analysis was performed based on the large terminase subunits. Specifically, for each group of viral genomes of interest, their large terminase subunits were first aligned by MUSCLE v5.1 (ref. ^47^) with default parameters. Then, we built phylogenetic trees based on the alignment results using FastTree v2.1.11 (ref. ^48^) with default parameters. Finally, iTOL^49^ was used to visualize the resultant trees.

